# Tree diversity effects on forest productivity increase through time because of spatial partitioning

**DOI:** 10.1101/2020.02.20.958553

**Authors:** Shinichi Tatsumi

## Abstract

**Background:** Experimental manipulations of tree diversity have often found overyielding in mixed-species plantations. While most experiments are still in the early stages of stand development, the impacts of tree diversity are expected to accumulate over time. Here, I present findings from a 31-year-old tree diversity experiment (as of 2018) in Japan.

**Results:** I find that the net diversity effect on stand biomass increased linearly through time. The species mixture achieved 64% greater biomass than the average monoculture biomass 31 years after planting. The complementarity effect was positive and increased exponentially with time. The selection effect was negative and decreased exponentially with time. In the early stages (≤3 years), the positive complementarity effect was explained by enhanced growths of early- and mid-successional species in the mixture. Later on (≥15 years), it was explained by their increased survival rates owing to vertical spatial partitioning — i.e., alleviation of self-thinning via canopy stratification. The negative selection effect resulted from suppressed growths of late-successional species in the bottom layer.

**Conclusions:** The experiment provides pioneering evidence that the positive impacts of diversity-driven spatial partitioning on forest biomass can accumulate over multiple decades. The results indicate that forest biomass production and carbon sequestration can be enhanced by multispecies afforestation strategies.

## Background

The positive relationship between biodiversity and ecosystem functioning (BEF) has been widely observed in experimental manipulations (Cardinale et al. 2007; Tilman et al. 2014). Species mixtures often achieve higher performance, typically measured by biomass productivity, than expected from average monoculture performance. This net biodiversity effect can be additively partitioned into complementarity and selection effects (Loreau and Hector 2001). The complementarity effect quantifies the average species overyielding which typically derive from niche partitioning, facilitation, and/or ecosystem feedbacks (Eisenhauer et al. 2012; Reich et al. 2012). Selection effect arises when species that produce large biomass in monocultures tend to overyield or underyield in mixtures (i.e., positive or negative selection) (Loreau and Hector 2001). Results from grassland diversity experiments have shown that the complementarity effect often becomes increasingly important with time (Fargione et al. 2007; Cardinale et al. 2007; Reich et al. 2012).

In the last two decades, diversity experiments using trees have been established to advance BEF research and to inform the increasing afforestation practices worldwide (Verheyen et al. 2016; Paquette et al. 2018). Compared to grasslands, forests develop over long time scales and form complex spatial structure. The development and complexity of stand structure are often largely driven by interspecific variation in successional niches (*sensu* Pacala and Rees 1998). Successional niches of tree species are typically represented by rapid growth under ample resources vs. survival in deep shade (Wright et al. 2010). While most tree diversity experiments are still in the early stages of stand development (≤20 years, mostly ∼10 years), variation in successional niches is expected to become relevant as stands enter canopy closure and the self-thinning stages (Grossman et al. 2018).

Here, I present findings from a 31-year-old tree diversity experiment (as of 2018) in northern Japan. The experiment was set up in 1987 and, to my knowledge, it is the oldest running diversity experiment in the world. By 2018, forest canopies had closed and some trees had reached >20 m in height. I expected the diversity effect on forest biomass to increase through time along with stand development. Specifically, I expected to see increasing complementarity effect as species mixing allowed spatial partitioning among species with different successional niches.

## Methods

### Experimental design

The experiment was set up within the tree nursery of the University of Tokyo’s Hokkaido Forest in Furano, northern Japan (43°13 09” N; 142°22’53” E; 223 m elevation). The dominant natural forest type in the region is cool-temperate conifer–broadleaf mixed forest. The mean annual temperature is 6.3 °C. The mean monthly temperatures range from a minimum of −7.2 °C in January to a maximum of 21.5 °C in August (University of Tokyo Forests, 2018). The average annual precipitation is 1,210 mm. The soil type is dark-brown forest soil. I refer to the experiment as the Furano experiment based on the name of the region.

In 1987, three native tree species were planted: *Betula maximowicziana* (Monarch birch), *Quercus crispula* (Japanese oak), and *Abies sachalinensis* (Todo fir). These species are often considered as dominant early-, mid-, and late-successional species in the region, respectively. The first two species are deciduous broad-leaved species and the last species is an evergreen conifer species. One monoculture for each of the three species and one mixture of the three species were established. Plot sizes were 6 m × 8 m (48 m^2^) for monocultures and 10 m × 10 m (100 m^2^) for the mixture. The plots were approximately 1 m apart from each other. The plots were relatively small in size and close to each other due to limited availability of land with uniform environmental conditions in the study site. The plot size of the mixture was set larger than that of the monocultures to equalize the numbers of individuals per tree species between the treatments as much as possible. Within each plot, trees were laid out hexagonally with 50-cm spacing, resulting in a density of 48,000 trees·ha^−1^. In the mixture plot, trees were planted such that their closest neighbors were always heterospecific (Fig. 1). The initial heights of all trees were set equal to ca. 35 cm by using 1-, 2, and 6-year-old samplings for *B. maximowicziana, Q. crispula*, and *A. sachalinensis*, respectively. The entire experiment consisted of plots with other treatments as well (namely, tree densities and species combinations; Moriyama et al. 2004), but these plots could not be used in this study because the trees were not measured before 2018.

**Fig. 1.**
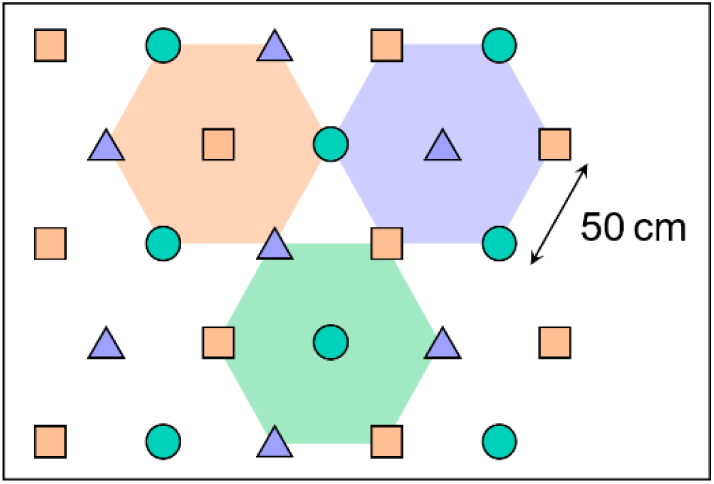
Planting layouts in the mixture plot of the Furano diversity experiment. Different symbols indicate different tree species. Trees were planted such that their closest neighbors are always heterospecific, as indicated by the hexagons.

### Tree measurements

Tree sizes were measured in 1988, 1989, 1990, 2002, 2003, and 2018 (=1, 2, 3, 15,16, and 31 years after planting). The heights of all live trees and diameters at breast height (DBH) of individuals with height ≥1.3 m were measured. Tree survival (i.e., alive or dead) of all individuals were additionally recorded in 2009 (=22 years after); i.e., 1988, 1989, 1990, 2002, 2003, 2009, and 2018. The aboveground biomass of each individual was estimated using an allometric equation in which the biomass was expressed as a function of either height or DBH (see Supplementary Materials for detail). In some measurement years, the height or DBH of some individuals were not measured. In such cases, the values were estimated using allometric equations (see Supplementary Materials for detail). The trees within 2 m of the plot edges were excluded from analyses to minimize the potential influences of edge effects. Fifty-one trees were planted in each monoculture and 175 trees were planted in the mixture (56 trees for each *B. maximowicziana* and *Q. crispula*, and 63 trees for *A. sachalinensis* — the difference among species derived simply from the spatial orders of trees within the plot), excluding those in the edge area. The entire experimental area was fenced to exclude large mammals.

### Statistical analyses

The net diversity effect (Δ*Y*) was defined as the difference between the total aboveground stand biomass (AGB), or yield, of the mixture and the average AGB of the monocultures (Loreau and Hector 2001). I estimated its additive components, complementarity and selection effects, according to Loreau and Hector (2001):

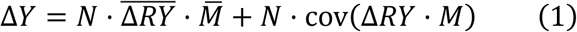

where *N* is species richness in the mixture (= 3), Δ*RY* is the deviation from the expected relative AGB for each species in the mixture, and *M* is the observed monoculture AGB. In calculating Δ*RY*, I quantified the observed relative AGB of each species as *Y*/*M*, where *Y* is the observed mixture AGB of the given species. The expected relative AGB was simply the inverse of species richness in the mixture (= 1/3) since species were planted in equal proportion.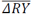 and 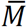 indicate the values averaged across all the species. The first and second components of Eq. 1 represent the complementarity and selection effects, respectively.

The temporal changes in net diversity, complementarity, and selection effects (*Y*) across the experiment year (*X*) were analyzed by fitting linear models *Y* = α + β · *X* and exponential models *Y* = α · *X*^β^ to the data. I used exponential models because *Y* can accumulate over time and thus change non-linearly (Reich et al. 2012). The parameters were estimated using the least-squares method (which implicitly assumes normally-distributed response variables). I also calculated the 95% confidence interval for each model; i.e., the range within which the fitted model was expected to lie 95% of the time, if the data collections and analyses were to be repeated multiple times. Akaike information criterions (AIC) (see Supplementary Materials for the definition) were compared between the two models. To account for the potential temporal autocorrelations in *Y*, I tested whether *Y* increased or decreased more largely than expected from random walks across time by using Bayesian state space models (see Supplementary Materials for detail).

Tree survival rates were compared among treatments and species in each year by using generalized linear models. Bernoulli distribution was used to describe the binary responses (alive vs. dead) of individual trees. The models included the treatment (monoculture vs. mixture) and species identity as explanatory variables, and were analyzed for each year separately. The interaction term was not included because the models for some years did not converge. The sample size was equal to the total number of trees planted (*n*=328) for all years except the first two years in which 21 trees were not measured (thus *n*=307) (see Table S1). Tree heights among treatments and species in each year were analyzed by means of two-way ANOVA followed by Tukey’s multiple comparison tests. The treatment, species identity, and their interaction were included as explanatory variables. See Table S1 for sample sizes. All statistical analyses were conducted using R 3.5.2 (R Core Team 2018).

## Results

The net diversity effect and complementarity effect showed positive values and tended to increase across the experiment year (Fig. 2a). The selection effect showed negative values after the first year and decreased thereafter (Fig. 2a). The linear model had a lower AIC than the exponential model for the net diversity effect (*R*^2^ = 0.91), while the exponential models were selected over the linear models for the complementarity (*R*^2^ = 0.99) and selection effects (*R*^2^ = 1.00) (Table S2). State space models accounting for temporal autocorrelations also showed that the net diversity and complementarity effects increased over time while the selection effect decreased in negative direction (Table S3). The survival rates of individual trees did not differ significantly between the monoculture and mixture plots during the first three years (Fig. 2b, Table S4). From the 15th year onwards, however, trees in the mixture showed higher survival rates than those of the same species in the monocultures (Fig. 2b, Table S4). Interspecific differences in survival rates became significant from the second year onwards (Table S4). The survival rates of *B. maximowicziana* and *Q. crispula* were relatively low (Fig. 2b). After 31 years, only one large individual (21.2 m in height) persisted in the *B. maximowicziana* monoculture (Figs. 2b, 3b; Table S1). Trees that were shorter in height tended to show lower survival rates than taller trees of the same species (Fig. S1).

**Fig. 2.**
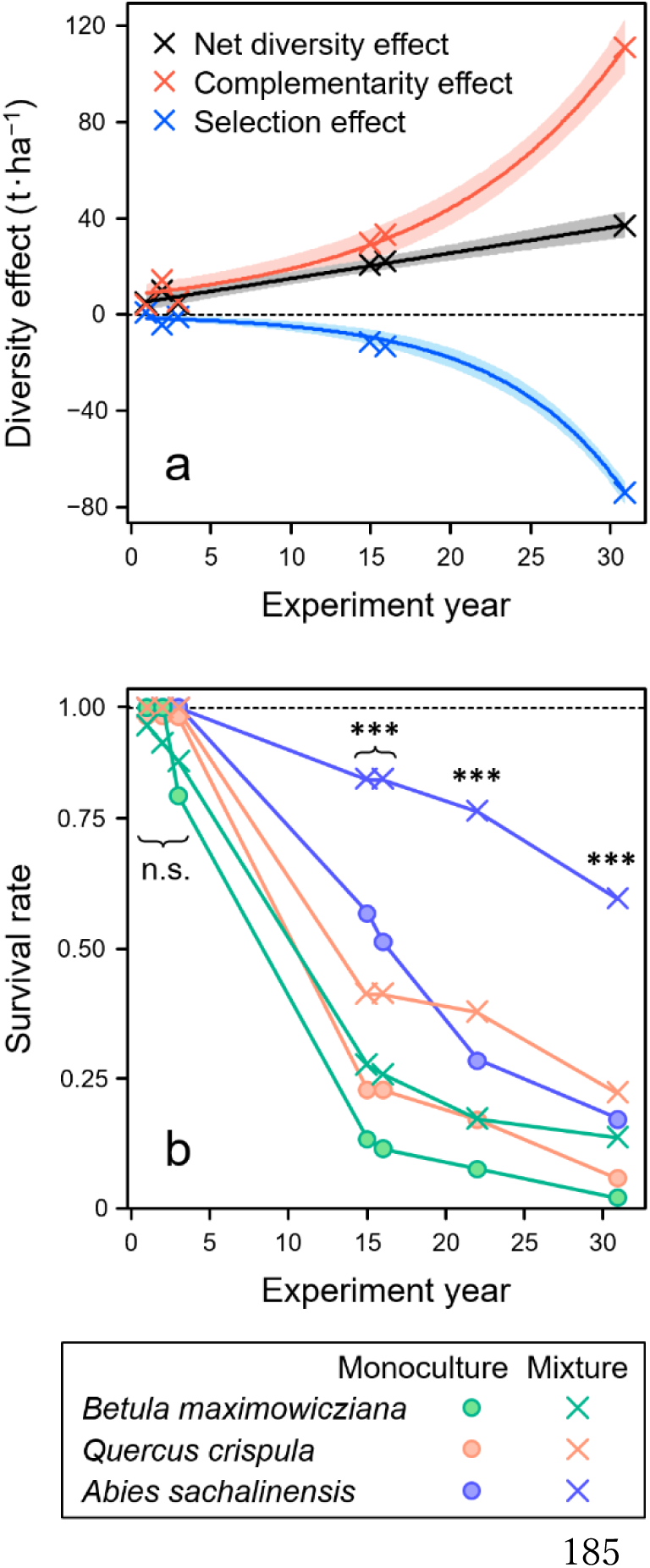
(a) Changes over time in net diversity effect, and its additive components, complementarity and selection effects, on aboveground stand biomass (t·ha^−1^). The crosses indicate observed values. The lines indicate fitted models, where linear model was selected for the net diversity effect and exponential models for the complementarity and selection effects. The shaded areas show the 95% confidence intervals of the fitted models. (b) Survival rates of three tree species in the monoculture and mixture plots. The circles and crosses indicate the observed values, calculated as *L*/(*L*+*D*), where *L* and *D* are the numbers of live and dead trees, respectively. Asterisks indicate significant differences in survival rates among treatments (monoculture vs. mixture) in each measurement year, as tested by generalized linear models with ‘treatment’ and ‘species identity’ as explanatory variables (see Table S4 for the results on species identity): ***, *P* < 0.001; n.s., *P* ≥ 0.05.

Three years after the onset of the experiment, *B. maximowicziana* and *Q. crispula* monocultures showed larger AGB than the average monoculture AGB (Fig. 3a). While *B. maximowicziana* showed the largest AGB among the monocultures, it accounted for a relatively small proportion of the total AGB in the mixture (Fig. 3a). This explains the negative selection effect in the third year (Fig. 2a). Fifteen years after planting, *A. sachalinensis* showed larger AGB than the average monocultue AGB, but constituted for the smallest proportion in the mixture (Fig. 3a). This discrepancy became even more pronounced 31 years after planting, explaining the escalating, negative selection effect aross time (Fig. 3a). On the other hand, *B. maximowicziana* and *Q. crispula* monocultures showed relatively low AGB after 15 years, and this trend became even clearer after 31 years (Fig. 3a). Their relative AGB were, however, comparatively higher in the mixture than in the monoculture (Fig. 3a). This explains the increasing complementarity effect through time (Fig. 2a).

**Fig. 3.**
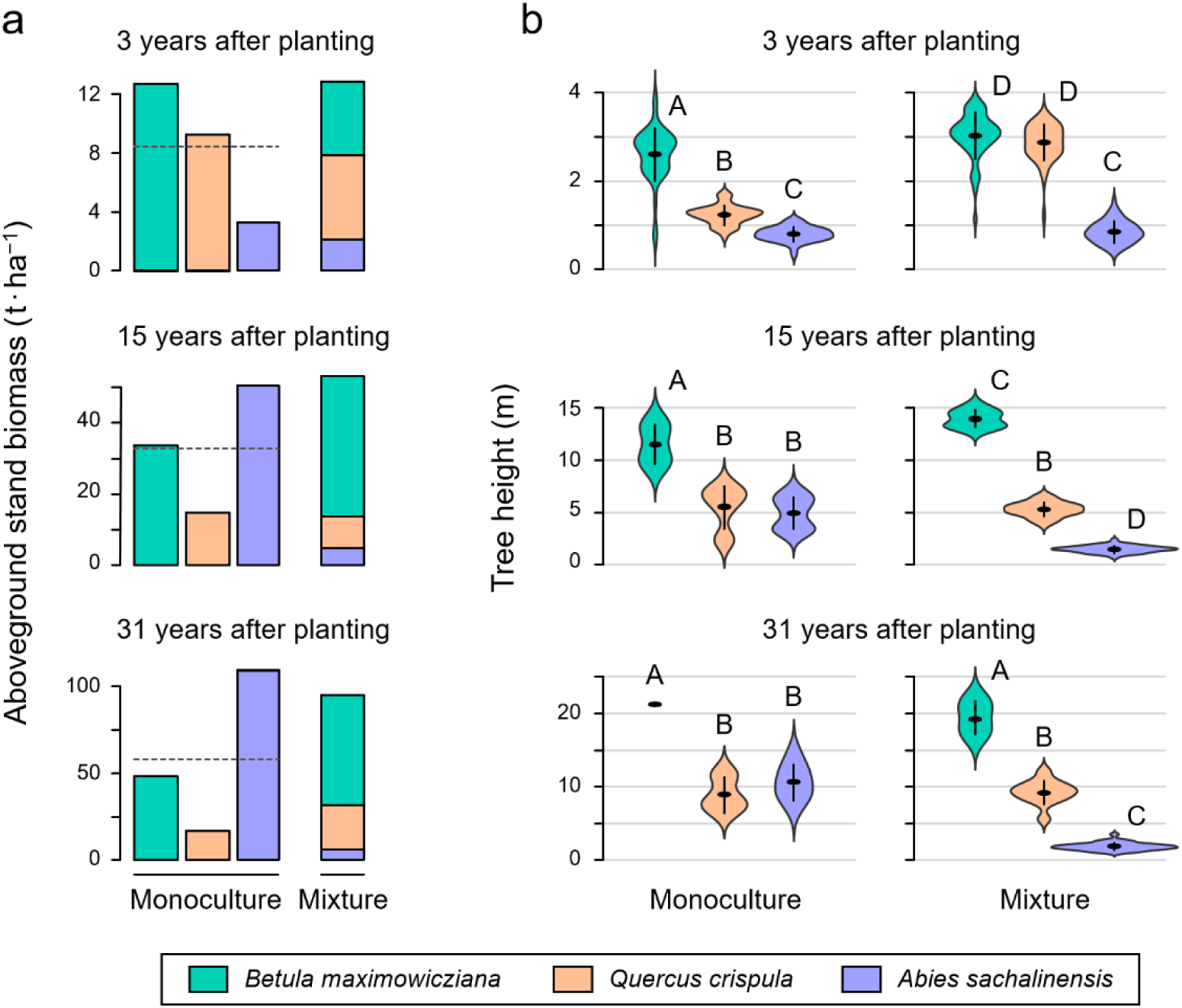
(a) Aboveground stand biomass in the monoculture and mixture plots in different years after planting. The horizontal dashed lines indicate the mean values across the monocultures. (b) Frequency distributions of tree height shown by violin plots. The horizontal and vertical lines indicate the mean values and the standard deviations, respectively. Different letters indicate significant differences (*P* < 0.05) among treatments and tree species in each measurement year, as tested by Tukey’s multiple comparisons following ANOVA.

Two-way ANOVA and Tukey’s tests revealed that *Betula maximowicziana* and *Q. crispula* in the mixture grew taller than those in the monocultures three years after planting (Fig. 3b; Table S5). After 15 years, tree heights in the mixture varied significantly among species, with *B. maximowicziana* being heigher and *A. sachalinensis* being shorter in the mixture than in the monocultures (Fig. 3b). Such vertical stratifications in the mixture were also observed 31 years after planting, where *B. maximowicziana, Q. crispula*, and *A. sachalinensis* occupied the height layers of ca. 20 m, 10m, and 2 m, respectively (Fig. 3b).

## Discussion

I found that the net diversity effect on stand biomass increased over time (Fig. 2a). Despite recent social concerns on mixture stands (Verheyen et al. 2016; Paquette et al. 2018), there has yet been a limited number of experimental investigations regarding the benefits of species mixing in time scales relevant to actual forestry and afforestation operations. Experimental studies using young tree stands have often found overyielding in species mixtures (Williams et al. 2017, Van de Peer et al. 2018; but see Grossiord et al. 2013; reviewed in Grossman et al. 2018). One of such studies also showed that the tree diversity effect increased over the first eight years of experiment (Huang et al. 2018). Adding on to these earlier findings, the Furano experiment showed that the species mixture achieved 64% greater biomass than the monocultures 31 years after planting (Fig. 3a). It also revealed that the complementarity effect increased more largely than has the selection effect decreased in negative direction with time (Fig. 2a). These results indicate that the impacts of species mixing can accumulate over time at least in some forest ecosystems, encouraging multispecies afforestation strategies to enhance long-term biomass production and carbon sequestration.

During the first three years of the experiment, early-and mid-successional species (*B. maximowicziana* and *Q. crispula*) grew significantly larger in the mixture than in the monocultures (Fig. 3b), a result which was attributable to increased availability of canopy space (Williams et al. 2017). The survival rates, on the other hand, did not differ significantly between the mixture and monocultures (Fig. 2b). These results coincide with previous findings in young tree stands that enhanced growth, but not survival rate, often contribute to overyielding (Huang et al. 2018; reviewed in Grossman et al. 2018). Later on the experiment (≥15 years), the cause of overyielding shifted from tree growth to survival. Trees in the mixture showed higher survival rates than those of the same species in the monocultures (Fig. 2b). This can be potentially explained by the reduction in competition-induced mortality owing to vertical partitioning — i.e., alleviation of self-thinning via canopy stratification (Fig. 3b). Note here that the vertical partitioning *per se* unlikely resulted from overyielding, but rather from the inherent differences in species’ successional niches, namely maximum growth rates under sufficient resources. I also found that the complementarity effect became increasingly large with time (Fig. 2a). This was likely because the intraspecific competition in each canopy layer intensified as crowns enlarged, and this impact was especially strong in the monocultures that had a single layered canopy (Fig. 3b).

The late-successional species (*A. sachalinensis*) achieved the highest monoculture biomass after ≥15 years (Fig. 3a). By contrast, in the mixture, its total biomass remained significantly small (Fig. 3a), which led the selection effect to take negative values (Fig. 2a). The low total biomass of *A. sachalinensis* was attributable to their suppressed growths (Fig. 3b), and was not explained by their survival rate, which was higher in the mixture than in the monoculture (Fig. 2b). In a grasslahnd experiment, Polley et al. (2003) found a negative selection effect when productive species were overtopped by less productive, early-growing species. Similarly, the suppressed growths of *A. sachalinensis* trees in the mixture (Fig. 3) was likely due to resource pre-emptions by species in the upper-layers (i.e., asymmetric competition for light; Weiner 1990). This view is also supported by earlier findings that, as a shade-tolerant species, *A. sachalinensis* can survive, but not necessarily grow, under low light levels (Iijima et al. 2009). I also found that the selection effect decreased over time (Fig. 2a). In grassland experiments, decreases in selection effect, as well as increases in complementarity effect, have shown to be driven by mechanisms such as nitrogen accumulations and biotic feedbacks (Fargione et al. 2007, Eisenhauer et al. 2012, Reich et al. 2012). It would be interesting to investigate how relevant these mechanisms are to forest ecosystems in the future.

Tree mortality is typically caused not only by competitive interactions but also ecological disturbance such as windstorms and insect outbreaks. In fact, the study region has experienced multiple windstorms over the past decades and an outbreak of herbivorous insects (Japanese giant silkworms; *Caligula japonica*) that fed on *B. maximowicziana* during 2006–2013 (= 19–26 years after planting). Such disturbance events could explain the low survival rates of *B. maximowicziana* and *Q. crispula* (Fig. 2b) to some extent. It is also possible that the positive impacts of species mixing on forest biomass growth (Fig. 1a) derived partly from a herbivory-mediated phenomenon known as associational effect (i.e., the interdependence of tree species with respect to their herbivores) (Cook-Patton et al. 2014). However, in the study region, herbivory-induced mortality is often contingent upon the levels of competitive stress to which individual trees had been subjected before the herbivory (Ohno et al. 2010). Moreover, trees that were shorter in height tended to show lower survival rates than taller trees of the same species (Fig. S1). Taken together, it is likely that disturbance played minor roles compared to light competition in the Furano experiment. I should, however, stress that this interpretation requires future verifications, given the fact that the experiment lacks plot replication and disturbance-related data.

## Conclusions

The Furano experiment provides the first experimental evidence that the tree diversity effect on biomass productivity can accumulate over multiple decades. The spatial partitioning induced by successional niche differences led to competition reductions in the upper layers (complementarity effect) and growth suppressions in the bottom layer (negative selection effect). The causes of overyielding also shifted over time, from enhanced tree growths in the mixture during the early stages (≤3 years) to increased survival rates later on (≥15 years). It should be noted, however, that the experiment has limited numbers of plots and treatments. Thus, the generality of the findings presented here should be examined carefully in future. Perhaps in combination with other tree diversity experiments (e.g., TreeDivNet; http://www.treedivnet.ugent.be), the data of the Furano experiment can be further utilized. The Furano experiment is unique in that it was set up in 1987, prior to the dawn of modern research on BEF (Vitousek and Hooper 1993). It is expected that the experiment will enter the next stage of stand development in the coming decades, where the early-successional species would die out and the remaining species take over the canopy space. The experiment would continue to provide, together with other tree diversity experiments, irreplaceable opportunities for long-term BEF research in forests.

## Acknowledgements

I thank Sadamoto Watanabe, who established the experiment, and Kiyoshi Ishida, Teruhisa Moriyama, and Takashi Masaki for providing field data collected before 2018. I am grateful to Naoto Kamata, Ayuko Ohkawa, Noriyuki Kimura, Satoshi Fukuoka, Masae Ando, and Hiroshi Inukai for their assistance in the fieldwork in 2018. I thank Adriano Roberto and Luke Potgieter for English language editing.

## Availability of data and materials

The datasets used in this study are available from the University of Tokyo’s Hokkaido Forest or the author upon request.

## Author’s contributions

ST conceived the idea, led the fieldwork in 2018, analyzed the data, and wrote the manuscript.

