## SupplementaryMaterials for "Tree diversity effects on forest productivity increase through time because of spatial partitioning"

### Supplementary Materials

#### Allometric equations

To estimate the aboveground biomass of the individual trees, I used the following allometric equations which were parameterized using *Betula ermanii*, *Quercus crispula*, and *Abies sachalinensis* in Hokkaido (Takagi et al. 2010, Ohtsu et al. 2015):

$$\ln(AGB) = 2.833 \cdot \ln(H) - 1.184 \qquad (1)$$

$$\ln(AGB) = 2.248 \cdot \ln(DBH) - 2.282 \qquad (2)$$

where $AGB$ is the total dry aboveground biomass of the stem, branches, and leaves (kg), $DBH$ is the diameter at breast height (cm), and $H$ is tree height (m). I used the first equation (Ohtsu et al. 2015) for trees with $H$ <1.3 m and the second equation (Takagi et al. 2010) for trees with $H$ ≥1.3 m. To correct for the inevitable discrepancy in $AGB$ near $H$ = 1.3 m (i.e., $DBH \fallingdotseq 0$ cm) between the equations, I assumed $AGB$ = 0.643 kg for all trees with $H$ ≥1.3 m and $DBH$ ≤2.1 cm (i.e., where the $AGB$ estimates of the two equations matched).

From 1988 to 1990, some of the *Betula maximowicziana* and *Q. crispula* trees in the experimental site had reached 1.3 m in height, but their $DBH$ were not measured. I therefore estimated the $DBH$ of *B. maximowicziana* and *Q. crispula* trees using the following equations 3 and 4, respectively:

$$DBH = 0.127 \times \exp(0.949 \cdot H) \qquad (3)$$

$$DBH = 0.285 \times \exp(0.658 \cdot H) \qquad (4)$$

where $DBH$ and $H$ were defined the same as above. The equations 3 and 4 were estimated using *B. maximowicziana* and *Q. crispula* trees, respectively, which were planted in a different plot within the experimental site. In this plot, *B. maximowicziana, Q. crispula*, and *A. sachalinensis* were planted 1 m apart from each other. The trees were planted at the same time with trees in the other plots (i.e., 1987). The $DBH$ and $H$ in 1990 were used for the estimation. The sample size and $R^2$ were *n* = 29 and $R^2$ = 0.905 for *B. maximowicziana* and *n* = 42 and $R^2$ = 0.677 for *Q. crispula*.

In 2002 and 2003, the heights of trees with $DBH$ <3.5 cm were not measured in the *A. sachalinensis* monoculture. I therefore estimated their heights using the following equation:

$$H = 1.3 + 0.816 \cdot DBH \qquad (5)$$

where $DBH$ and $H$ were defined the same as above. The parameters were estimated using the $DBH$ and $H$ of trees with $DBH$ ≥3.5 cm in the same monoculture and *A. sachalinensis* trees planted in the 1 m-spaced mixed stand (*n* = 52; $R^2$ = 0.948).

#### State space modelling

I tested whether the net diversity, complementarity, and selection effects increased or decreased through time using Bayesian state space models:

$$Y_t \sim N(y_t, \sigma_\varepsilon{}^2)$$
$$y_t \sim N(y_{t-1} + \delta, \sigma_\eta{}^2) \qquad (6)$$

where $Y_t$ is the observed value and $y_t$ is the state variable of the net diversity, complementarity, or selection effect at time $t$, $\delta$ is the parameter that determines whether $y_t$ changed deterministically through time, and $\sigma_\varepsilon{}^2$ and $\sigma_\eta{}^2$ are variations. Time $t$ was represented by the year after planting. The $y_t$ at $t = 0$ was set 0. The parameters $\delta$, $\sigma_\varepsilon{}^2$, and $\sigma_\eta{}^2$ were assumed as non-informative priors: $\delta \sim N(0, 10^4)$, $\sigma_\varepsilon{}^2 \sim Unif(0, 10^4)$, and $\sigma_\eta{}^2 \sim Unif(0, 10^4)$.

Model 6 takes into account the temporal autocorrelations by assuming that $y_t$ is determined as $y_t = y_{t-1} + \delta + \varepsilon$, where $\varepsilon$ is a Gaussian noise determined by $\sigma_\eta{}^2$. The null hypothesis here is $y_t \sim N(y_{t-1}, \sigma_\eta{}^2)$ (i.e., random walk). The constant value $\delta$ tests whether $y_t$ increased or decreased more largely than expected from stochastic fluctuations.

The posterior distributions of the parameters were estimated using the Markov chain Monte Carlo (MCMC) method implemented by JAGS ver. 4.3.0 (Plummer 2003). I obtained posterior samples using three independent MCMC samplings, in each of which 1000 values were sampled at a 5-step interval after a burn-in period of 5000 MCMC steps. I judged that the MCMC calculation had converged when the $\widehat{R}$ values of all the parameters were <1.1 (Gelman et al. 2013).

**Akaike information criterion**

Akaike information criterion (AIC) of the regression models, for which the least-squares method was used to estimate the parameters, was defined as follows (Burnham and Anderson 2004):

$$\mathrm{AIC} = 2(p + 1) + n[\ln(2\pi \tfrac{S}{n}) + 1] \qquad (7)$$

where $p$ is the number of parameters, $S$ is the residual sum of squares, and $n$ is simple size.

**Table S1.** Numbers of live trees (= trees of which the heights were measured) in each year.

| Treatment | Species | Initial | 1 | 2 | 3 | 15 | 16 | 22 | 31 |
|---|---|---|---|---|---|---|---|---|---|
| Monoculture | *B. maximowicziana* | 51 | 30[†] | 30[†] | 42 | 7 | 6 | 4 | 1 |
| | *Q. crispula* | 51 | 50 | 50 | 50 | 12 | 12 | 9 | 3 |
| | *A. sachalinensis* | 51 | 51 | 51 | 51 | 30 | 27 | 15 | 9 |
| Mixutre | *B. maximowicziana* | 56 | 54 | 52 | 50 | 16 | 15 | 10 | 8 |
| | *Q. crispula* | 56 | 56 | 56 | 56 | 24 | 24 | 22 | 13 |
| | *A. sachalinensis* | 63 | 63 | 63 | 63 | 54 | 54 | 50 | 39 |
| Total | | 328 | 304[†] | 302[†] | 312 | 143 | 138 | 110 | 73 |

† In the *B. maximowicziana* monoculture, only 31 trees out of the 51 initially-planted trees were measured in the first and second year, likely due to limitations in the workforce.

**Table S2.** Results of regression analyses on the relationship between diversity effects $(Y)$ and experimental years $(X)$.

| Response variable $Y$ | Model | Estimates | | AIC[†] | $R^2$ |
|---|---|---|---|---|---|
| | | $\alpha$ | $\beta$ | | |
| Net diversity effect | $Y = \alpha + \beta \cdot X$ | 4.398* | 1.058*** | **113.3** | 0.974 |
| | $Y = \alpha \cdot X^{\beta}$ | 8.057* | $0.5\times10^{-4}$** | 120.5 | 0.913 |
| Complementarity effect | $Y = \alpha + \beta \cdot X$ | −3.811 | 3.242** | 135.3 | 0.899 |
| | $Y = \alpha \cdot X^{\beta}$ | 8.056** | $0.9\times10^{-4}$*** | **120.5** | 0.992 |
| Selection effect | $Y = \alpha + \beta \cdot X$ | 8.208 | −2.184* | 135.3 | 0.802 |
| | $Y = \alpha \cdot X^{\beta}$ | −1.357* | $1.3\times10^{-4}$*** | **110.2** | 0.997 |

† The values of Akaike information criterion (AIC) are shown in bold face for the model which had lower AIC than the other (i.e., linear vs. exponential).

* $P < 0.05$, ** $P < 0.01$, * $P < 0.001$

**Table S3.** Posterior distributions of Bayesian state space models on temporal changes in diversity effects $(Y)$.

| Variable $Y$ | Parameter $\delta$ mean | 2.5% credible interval | 97.5% credible interval |
|---|---|---|---|
| Net diversity effect | 1.154 | 1.119 | 1.190 |
| Complementarity effect | 3.416 | 3.381 | 3.451 |
| Selection effect | −2.262 | −2.297 | −2.228 |

**Table S4.** Results of generalized linear model analyses on differences in survival rates (for which Bernoulli distributions were used to describe 'alive vs. dead' responses) among treatments and tree species. The values show likelihood-ratio chi-squares.

| Experiment year | 1 | 2 | 3 | 15 | 16 | 22 | 31 |
|---|---|---|---|---|---|---|---|
| Treatment (mono vs. mix) | 3.2 | 5.0 | 2.6 | 18.7*** | 23.5*** | 38.5*** | 36.7*** |
| Species | 4.6 | 9.3** | 14.1*** | 45.4*** | 48.7*** | 50.2*** | 34.7*** |

** *P* < 0.01, * *P* < 0.001

**Table S5.** Results of two-way ANOVA on differences in tree heights (m) among treatments and tree species (DF = degrees of freedom, MS = mean squares, *F* = *F*-ratio).

| | 3 years after planting | | | 15 years after planting | | | 31 years after planting | | |
|---|---|---|---|---|---|---|---|---|---|
| | DF | MS | *F* | DF | MS | *F* | DF | MS | *F* |
| Treatment (mono vs. mix) | 1 | 36.7 | 50.7*** | 1 | 66.6 | 4.4* | 1 | 327.4 | 10.3** |
| Species | 2 | 17.6 | 24.4*** | 2 | 13.1 | 0.9 | 2 | 37.9 | 1.2 |
| Treatment × Species | 2 | 22.5 | 31.1*** | 2 | 129.7 | 8.5*** | 2 | 75.0 | 2.4 |
| Residuals | 306 | 0.7 | | 137 | 15.3 | | 65 | 31.9 | |

* *P* < 0.05, ** *P* < 0.01, * *P* < 0.001

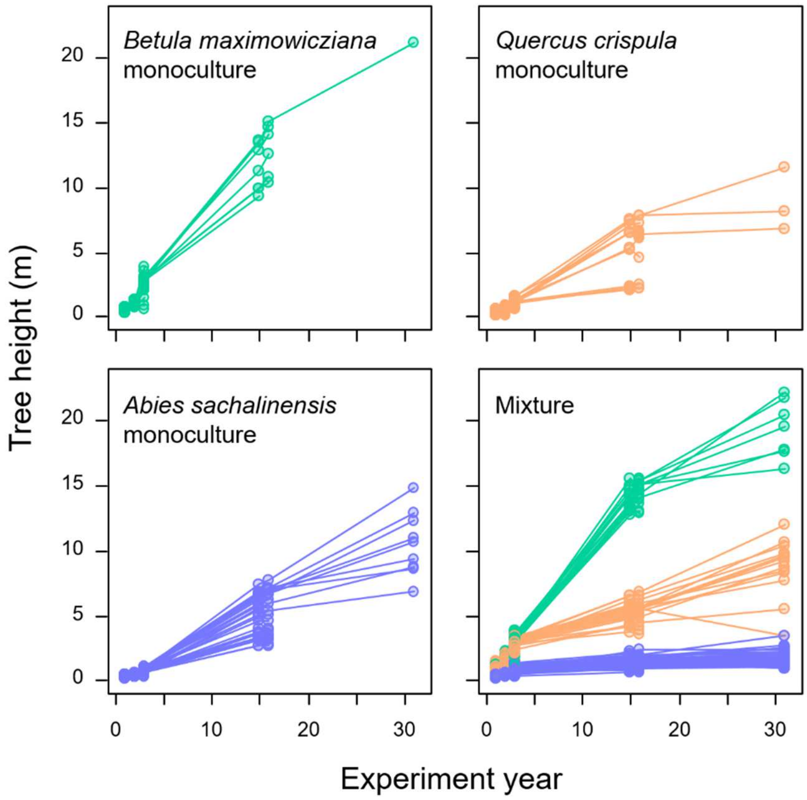


**Fig S1.** Changes in tree heights across experimental years in the Furano experiment. Each set of circles connected by lines represents the change in height of an individual tree. The lines are terminated when the trees were dead. The height of some trees decreased between some measurement intervals due to stem bending or breakage.
